# Massive presence of the invasive *Polydora websteri* (Polychaeta: Spionidae) in the North Adriatic Sea (Mediterranean Sea)

**DOI:** 10.1101/2024.07.18.603787

**Authors:** Marco Lezzi, Cristina Mazziotti

## Abstract

In recent decades, the Mediterranean Sea has experienced a marked escalation in the introduction of non-native and invasive marine species.

The global expansion of aquaculture, coupled with the increased movement of commercial species, has raised concerns about the potential for introducing parasites into new environments, posing a growing threat to marine ecosystems.

*Polydora websteri* is a shell boring polydorin worm that induces the formation of mud-filled blisters within the shells of molluscs. The annelid is considered indigenous to the Asian coasts and has been documented in various locations across the globe. In the present work, we report a massive presence of *P. websteri* in several sites of the North Adriatic costline and we provide some diagnostic features that are useful to identify the species. *Polydora websteri* can be distinguished of a lateral flange on chaetiger 5. Moreover the palps are characterized by continuous black lines along the margins of the ciliated groove.

The combined findings of this study provide compelling evidence that *P. websteri* is expanding its range within the Mediterranean coast. Given the potential for severe ecological and economic impacts associated with this invasive polychaete, this study serves as an urgent call to action for increased monitoring of *P. websteri* in both farmed and wild Pacific oysters in the Mediterranean Sea. Identification key of Mediterranean *Polydora* is provided.

## 1. Introduction

The expansion of global trade in recent decades has led to a significant increase in the introduction of species worldwide.

In addition to the transportation of species through ballast water or as hull fouling on ships, the deliberate introduction of commercially valuable species and the subsequent movement of farmed specimens within aquaculture sites are the primary pathways by which marine ecosystems are invaded (Katsanevakis et al. 2013). For example, the farming of marine molluscs has been recognized as a major route of species introduction, primarily due to the routine large-scale movement of stocks associated with this practice (Wolff and Reise 2002, Mckindsey et al. 2005, Ruesink et al. 2005).

Numerous species, including the Pacific oyster *Magallana gigas* (Thunberg, 1793), have been extensively translocated to novel ecosystems and are now globally traded, and cultured (Ruesink et al. 2005). As a result, the inadvertent introduction of organisms inhabiting the oysters, whether they are sessile and boring species or parasites, has taken place outside of their native environments (Elton 1958, Padilla et al. 2011, Goedknegt et al. 2016).

Example of organisms that are particularly vulnerable to accidental introduction along commercial molluscs comprises shell-boring polychaetes, with polydorins being the predominant subgroup within this category, commonly known as a “mud worms” or “mud blister worms” (Tan et al. 2023).

These worms are known to penetrate and create burrow mussel shell (Simon et al. 2015). During infestations, the shell becomes severely damaged by the burrowing activities of the worm, which diverts energy from growth to shell host’s repair, with deleterious effects on host fitness (Kent 1979, Leonart et al. 2003). Additionally, the impairment of shell integrity consequent to shell damage renders oysters increasingly vulnerable to predation from crabs and other predatory organisms (Buschbaum et al. 2007).

*Polydora websteri* Hartman in Loosanoff and Engle, 1943 is a frequently co-introduced polydorin species that is commonly found alongside cultured molluscs. This species forming mud-filled blisters inside the shells of molluscs. Mud worms typically undergo a pelagic larval stage, after which the larvae settle on the external side of a calcareous shell (Clements et al. 2018). Then they form a U-shaped burrow with a pair of exterior openings and during the growth these borings perforate the inner shell of the valve, eliciting a response from the host that results in a rigid layer of nacre being developed to isolate the burrow (Zottoli and Carriker 1974, Simon et al. 2015, Clements et al. 2018). The worm burrows beneath the thin, calcareous layer generated by its host; as this enclosed area becomes filled with detritus, mud, and worm feces: the “mud blister” (Haigler 1969, Zottoli and Carriker 1974). Blisters compromising the presentation of oysters served on the half-shell and if a blister is accidentally punctured while shucking oysters, the mud and feces can contaminate the oyster meat, making it unfit for consumption (Shinn et al. 2014). Mud blisters are a direct consequence of the feeding behavior of the polychaete, which results in the accumulation of detritus, mud, and fecal matter within the burrow. The host mollusc responds to this intrusion by producing additional layers of shell material to cover the burrow. The polychaete is widely recognized as a significant pest in commercial marine aquaculture worldwide, as the presence of mud blisters within the mollusc shells leads to a decrease in their market value (Kent 1979, Leonart et al. 2003, Shinn et al. 2014, Clements et al. 2018).

Historically, *P. websteri* and related polydorins have significantly undermined and even brought to a standstill oyster aquaculture industries worldwide. In the late 1800s, the unintentional introduction of the parasitic polychaete worm *P. websteri* alongside transplanted oysters played a pivotal role in the decimation of subtidal oyster beds across New South Wales, Australia (Whitelegge 1890, Ogburn et al. 2007). Similarly, the introduction of *P. websteri* via oyster transplants from Kaneohe Bay to Kakuku, Hawaii, resulted in widespread and severe damage to shellfish production (Bailey-Brock 1982). Oyster farms along the east coast of the United States have endured a persistent infestation of *P. websteri* since the 1940s, leading to significant losses in oyster aquaculture production (Lunz 1940, Loosanoff and Engle 1943). These examples serve as compelling evidence of the remarkable invasive potential of *P. websteri*, demonstrating its ability to successfully establish itself in new environments and, once established, inflict substantial damage on aquaculture production.

Research indicate that *P. websteri* is deemed to be native to Asian coasts (Rice et al. 2018) and has been reported from several locations all over the world, including Australia, New Zealand, Hawaii, Brazil, the Atlantic and Pacific coasts of North America, the Black Sea, Suez Canal and Wadden Sea in Europe (Lunz 1940, Loosanoff and Engle 1943, Read 2010, Surugiu 2012, Sato-Okoshi and Abe 2013, Barros et al. 2017, Ye et al. 2017, Rice et al. 2018, Abd-Elnaby 2019, Martinelli et al. 2020, Waser et al. 2020, Rodewald et al. 2020), and was more recently recognized by Mikac et al. (2023) in the Adriatic sea.

Genetic homogeneity between North American, Hawaiian, and Asian specimens also suggests that human-mediated transport produced high levels of connectivity, supporting the notion that it is native to Asian coasts (Rice et al. 2018, Sato-Okoshi and Abe 2013). In the present work, shells of Pacific oysters *M. gigas* were investigated for signs of polydorin polychaetes throughout the North Adriatic Sea. Our results confirm the a massive presence of *P. websteri* in several sites of the coast of Emilia Romagna (Italiy) in wild populations of *Magallana gigas*. This study describe the species, including key diagnostic features that distinguish it from similar species currently known in the Mediterranean Sea.

## 2. Materials and Methods

### Oyster collections

To assess whether shell-boring polychaetes were present in North Adriatic coast oysters (*Magallana gigas*) and to confirm the species identity of these worms, we collected about 30 grown oysters from the port of Ravenna, Italy (44°29’27.7”N 12°17’24.2”E) in March and August 2023 (Figure 1). In the habour of Goro (44°50’29.1”N 12°17’14.5”E) and Cesneatico (44°12’21.4”N 12°23’47.1”E) in June 2024). The oysters were collected manually from artificial substrates in a tourist harbour area.

**Figure 1.**
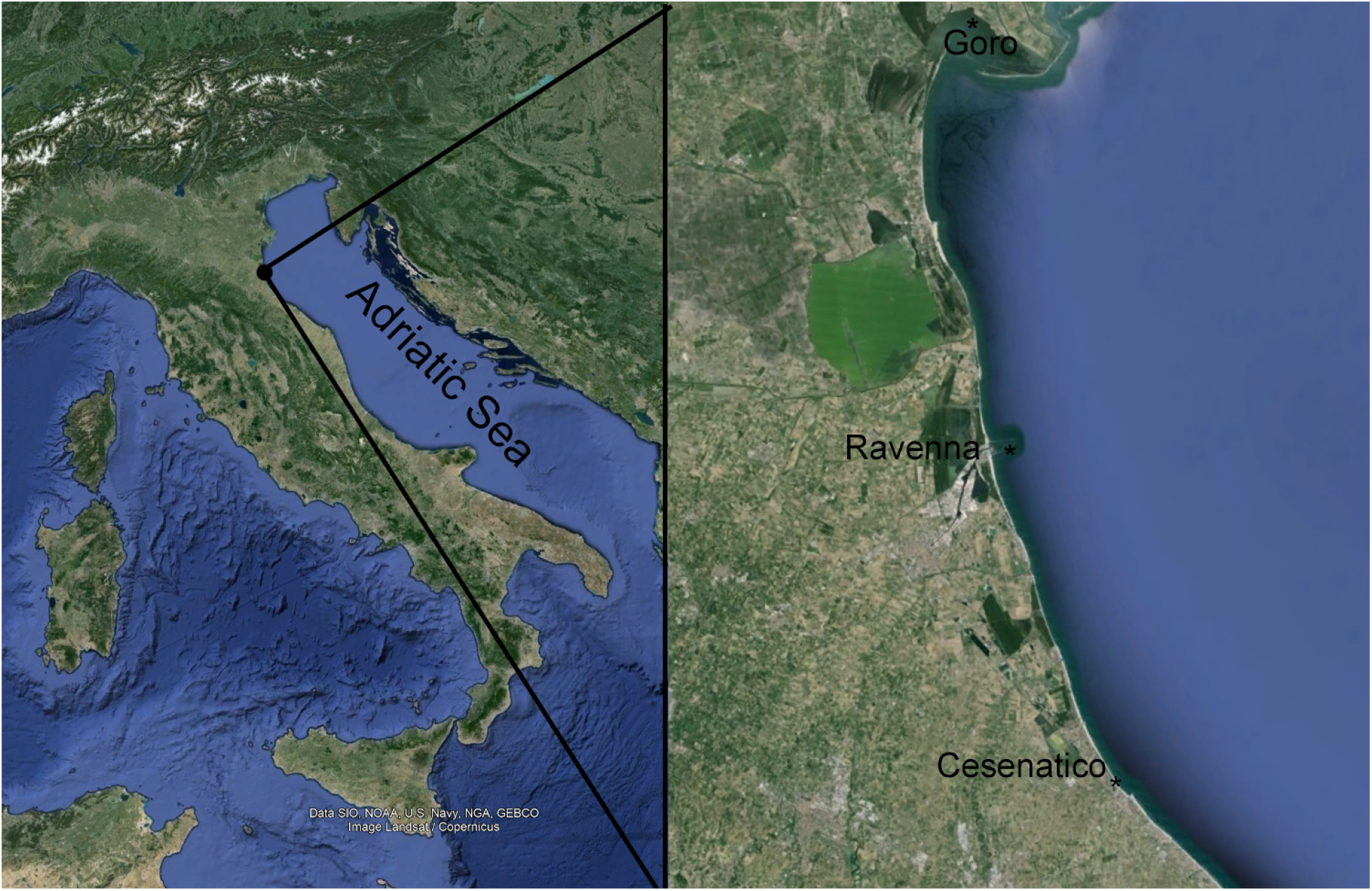
Sampling sites in the North Adriatic Sea:

### Worm collections

The oysters were shucked and the soft tissues were removed. To detect the presence of mudworm infestation, we examined the right and left valves of oysters under a stereomicroscope Nikon SMZ18, such as burrows and blisters (Figure 2 a, b, c). Mudworms were extracted manually from blisters or burrows using tweezers. For morphological examination, worms were preserved in 4% buffered formalin for two days and transferred to 70% ethyl alcohol (EtOH) for storage.

**Figure 1.**
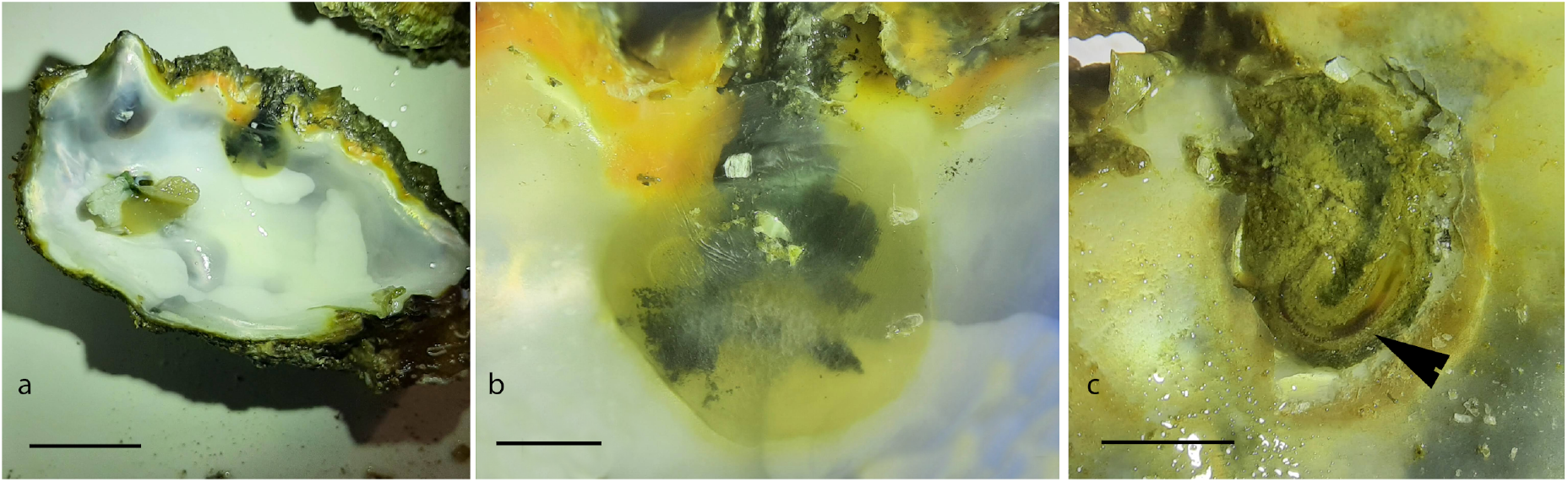
*Magallana gigas* infested with *Polydora websteri* collected from Ravenna. (**a**) Inner surface of an infested valves, (**b**) closed mud blister filled with mud, detritus, and feces (**c**) Opened mud blister, arrow indicates the mud blister with *P. websteri*. Scale bar: a, 3 cm; b, 1 cm; c, 1cm. Photo credit: Marco Lezzi.

The specimens of *P. websteri* from the Adriatic Sea were sent to the Turin Museum of Natural History (MRSN), and were also deposited in the Laboratory of Benthos Ecology collection of the Oceanographic Unit Daphne of ARPAE, Regional Agency for Environmental Protection and Energy of Emilia Romagna.

## 3. Results

In total, 40 Pacific oysters were observed systematically for *Polydora* spp. infestation. The prevalence was 70% in alive Pacific oysters with at least one blister or burrow per shell. In each oyster approximately 2 ± 0.5 polydorin were found and identified as *P. websteri*.

## *Polydora websteri* Hartman in Loosanoff and Engle, 1943

*Polydora websteri*:in Loosanoff and Engle, 1943

*Polydora websteri:* Loosanoff and Engle 1946: 70–72, Fig. 1; Blake 1969: 814–815, Fig. 2; Blake 1971: 6–8, Fig. 3; Foster 1971: 26; Blake et al. 1978: 258–259, Figs 43k–n ;Handley et al. 1997: 191–205; Radashevsky and Williams 1998: 212–216; Radashevsky 1999: 107–113, Fig. 1; Sato-Okoshi 1999: 832–834, Fig. 2B; Surugiu 2005: 67; Read 2010: 9–11, Figs 1H–J, 2B, 2D, 2F, 4D–G; Surugiu 2012: 50–53, Fig. 3; Sato-Okoshi and Abe 2013: 1280–1281, Fig. 2; Rice et al. 2018; Ye et al. 2017; Martinelli et al. 2020.

### Material examined

Italy, North Adriatic Sea, Ravenna Harbour, 4 specimens, March, August 2023, C. A. Lezzi, from wild *Magallana gigas*. Italy, North Adriatic Sea, Goro touristic Harbour, 1 specimens, May, 2024, C. A. Lezzi, from wild *Magallana gigas*. Italy, North Adriatic Sea, Cesenatico Harbour, 2 specimens, May 2023, C. A. Lezzi, from wild *Magallana gigas*.

### Description

Adult worms up to 15 mm long, 0.5 mm wide at chaetiger 5, and with up to 116 chaetigers. Colour in life light tan with red blood vessels in branchiae and palps. Preserved specimens pale yellowish. Middle-distal part of palps with a black sinuous line along edges of ciliated food groove, persisting in preserved specimens (Figure 3 a, b). Anterior margin of prostomium slightly incised, caruncle extending back to the end of chaetiger 2. Occipital antenna absent. Four small rounded eyes in trapezoidal arrangement (Figure 3 e). Chaetiger 1 with well-developed neuropodial postchaetal lamellae and a fascicle of capillary neurochaetae; small lamellae on notopodium, notochaetae absent. Chaetigers 2–4 and 6 with well-developed and flattened noto and neuropodial postchaetal lamellae and limbate capillary chaetae. Chaetiger 5 broader and longer than adjacent ones, bearing an oblique row of 5–6 thick, falcate spines alternating with pennoned companion chaetae. Major spines of chaetiger 5 falcate, with a subterminal flange on lateral side (Figure 3d, f). Chaetiger 5 with up to six superior dorsal limbate capillary chaetae arranged in vertical row and a bundle of shorter ventral limbate capillaries. Neuropodial bidentate hooded hooks from chaetiger 7, up to 11 in a vertical row in middle chaetigers, decreasing to 2–3 posteriorly. Hooded hooks with main fang at right/acute angle to shaft and with wide acute angle (∼40°) to smaller apical tooth; shaft with a prominent constriction and distinct manubrium (Figure 3c). Posterior notopodia with capillaries, without special notopodial spines. Branchiae flat, from chaetiger 7, reaching maximum length by chaetigers 10-12, continuing to near the posterior end, absent from last 10-15 chaetigers, in the posterior half of the body decreasing gradually in size. Pygidium thin, cup-shaped; anus at base of dorsal gap.

**Figure 3.**
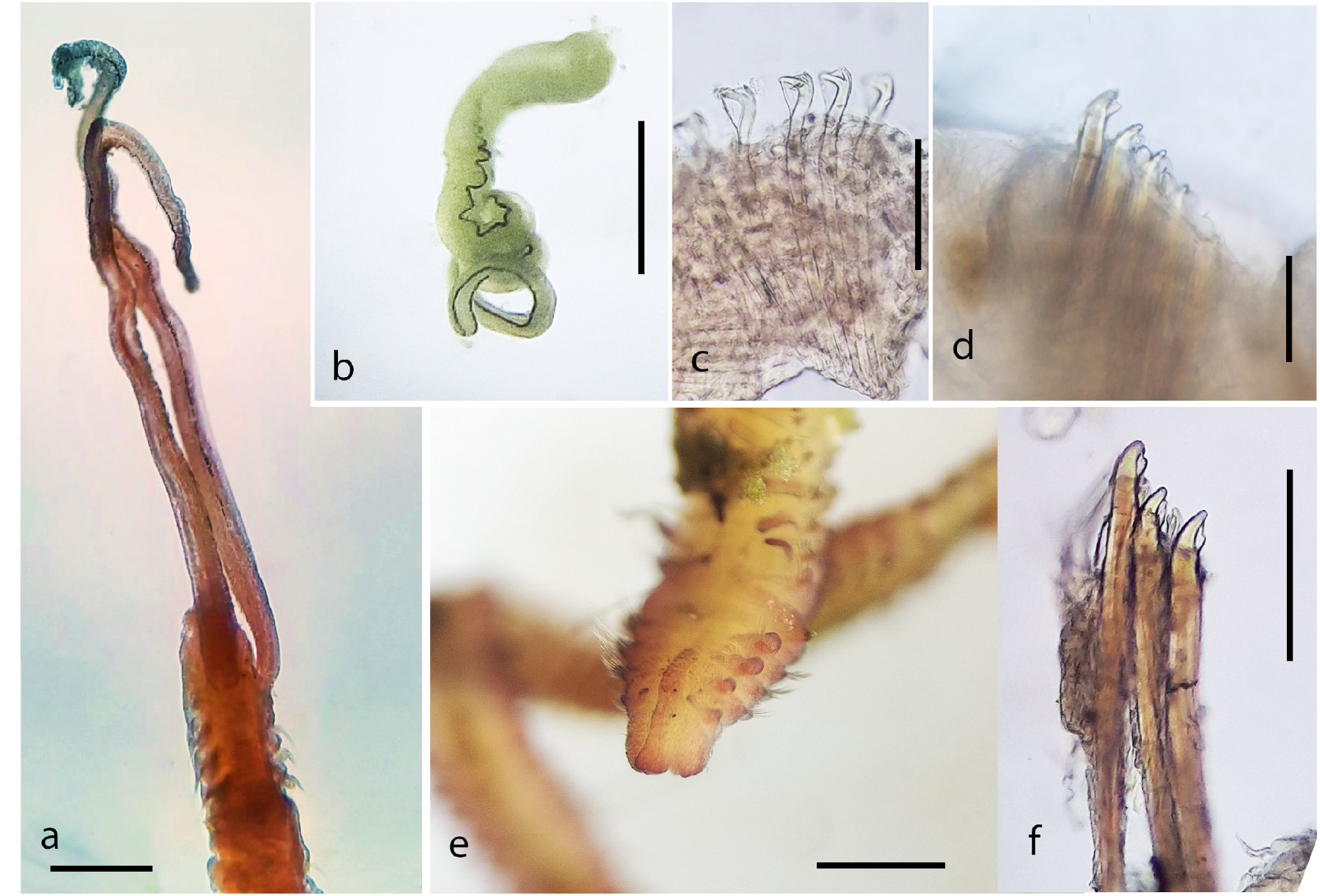
*Polydora websteri:* (**a**) Dorsal view of anterior end of specimen with palps attached; (**b**) palps; (**c**) neuropodial bidentate hooded hooks from chaetiger 12; (**d, f**) falcate spines of chaetiger 5 with a subterminal flange on concave side; (**e**) Dorsal view of anterior end of specimen stained with shilastain “A”. Scale bar: a, b, 600 µm; c, d, f, 50 µm; e 500 µm. Photo credit: Marco Lezzi.

### Remarks

The specimens agree with descriptions of the type material (Radashevsky et al. 1999) and descriptions of conspecifics found globally (e.g. Sato-Okoshi 1999, Surugiu 2012, Rice et al. 2018, Martinelli et al. 2020).

*Polydora websteri* is broadly similar in morphology to the Mediterranean *Polydora ciliata* (Johnston, 1838) and *Polydora limicola* Annenkova, 1934. However, the latter species differ from *P. websteri* in having a small lateral triangular tooth on the major modified spines of chaetiger 5 instead of a lateral flange.

*Polydora websteri* may be also confused with other species because on worn spines the flange may have the appearance of a flange and accessory tooth that are not connected, closely resembling the major spines of other *Polydora* species (i.e. *Polydora cornuta* Bosc, 1802) (Foster 1971, Çinar et al. 2005, Radashevsky 2005). Confusion could be avoided if deeper, unworn chaetae are examined. Moreover the palps of *P. websteri* are characterized by continuous black lines along the margins of the ciliated groove and this characteristic is distinctive of the identified species (Read et al. 2005).

*Polydora websteri* can be distinguished from *Polydora colonia* Moore, 1907 and *Polydora spongicola* Berkeley & Berkeley, 1950 because the latter have the spines of chaetiger 5 without a curved accessory tooth; *Polydora hoplura* Claparède, 1868 differs in having last chaetigers with recurved notopodial spines, while *Polydora agassizii* Claparède, 1869 differs mainly in having black spots on palps.

## Key for Mediterranean *Polydora* identification

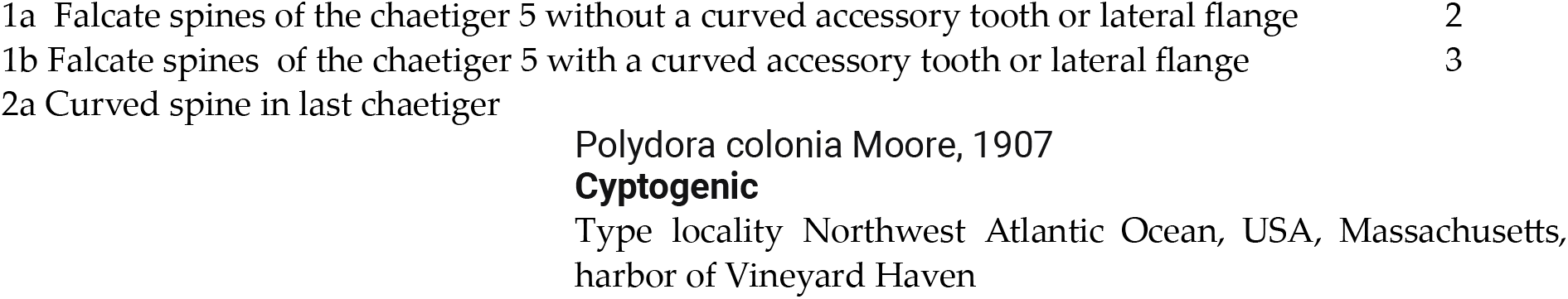
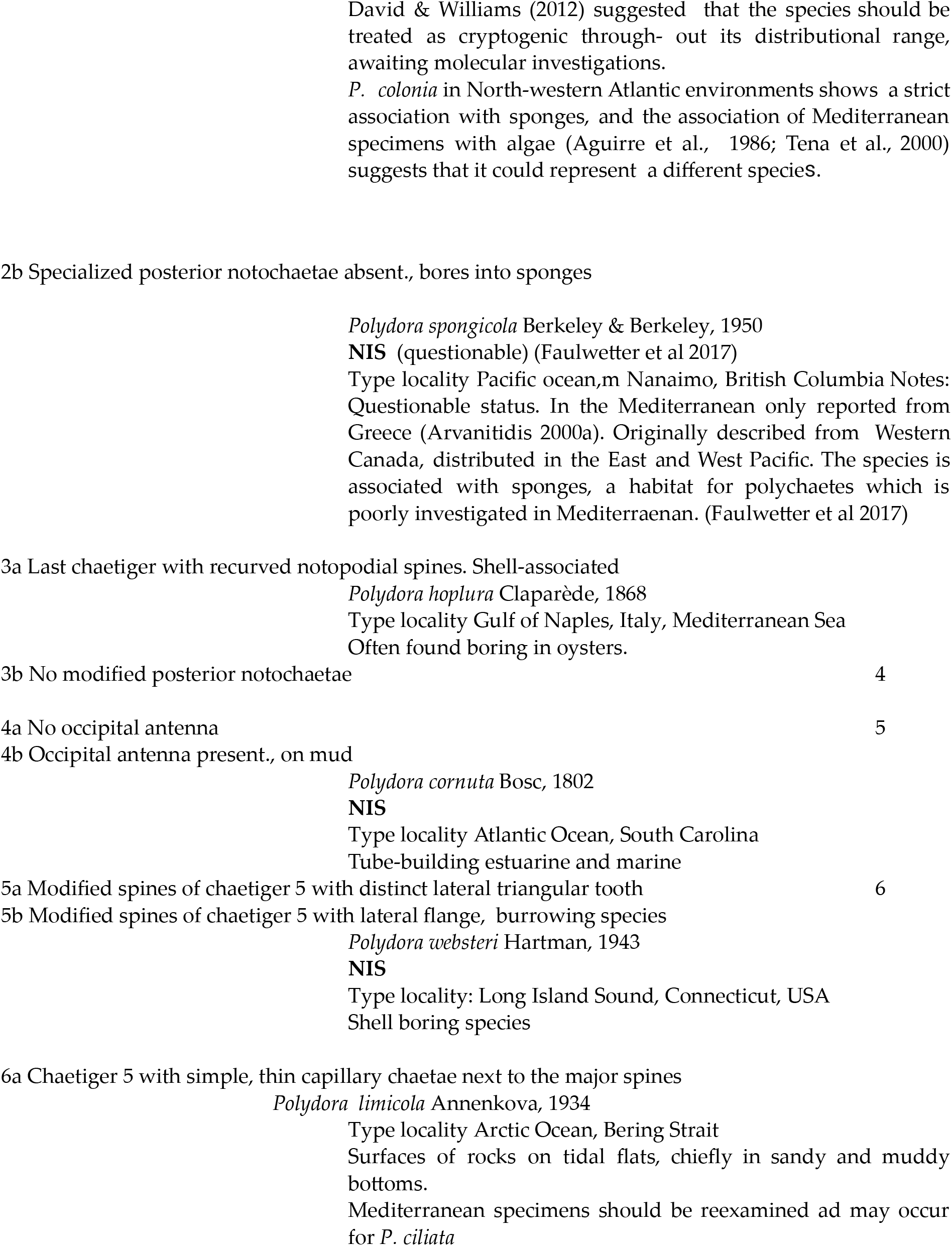
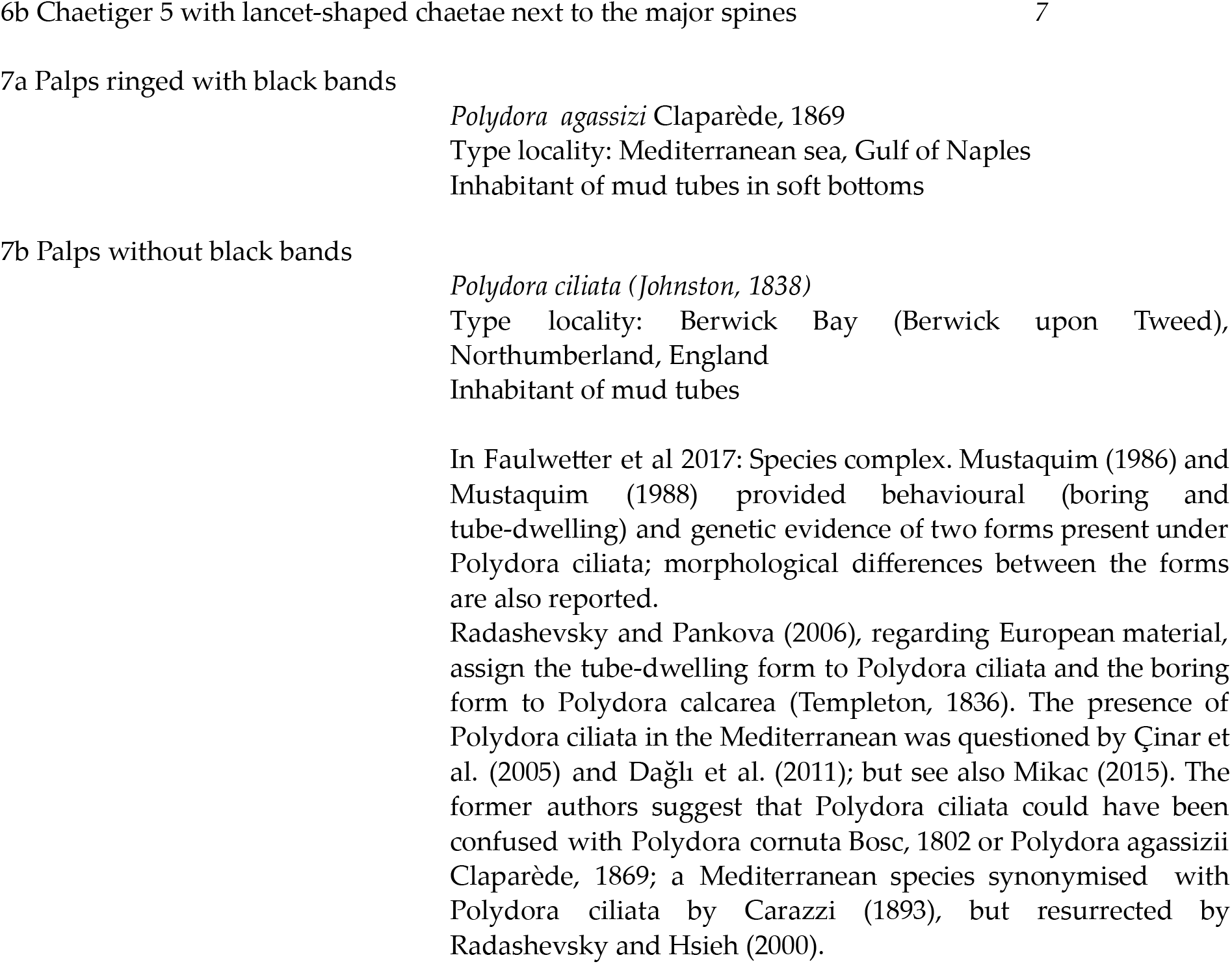

## 4. Discussion and conclusions

In the Mediterranean Sea were reported seven species of *Polydora* (Çinar et al. 2005, Coll et al. 2010, Bertasi 2016): *P. agassizii, P. ciliata, P. colonia, P. cornuta, P. hoplura, P. limicola, P. spongicola*. Among them, *P. cornuta* is considered an alien species (Çinar et al. 2005, Bertasi 2016) and the presence of the sponge-boring *P. colonia*, initially considered alien (see Occhipinti-Ambrogi et al. 2011), is now questionable in Italian waters (Langeneck et al. 2020).

The observation of *P. websteri* in several locations of the Emila Romagna coast shown in this study serves as a significant record after the one of Mikac et al. (2023).

At present, the species has invaded most of the seas and what was missing in its invasion history was the Mediterranean Sea.

Pathology caused by shell-boring mud worms such *P. websteri*, causes unsightly blisters; that significantly affects the aesthetic and market value of infested oysters, particularly those intended for consumption on the half-shell (Morse et al. 2015). The European oyster industry is largely driven by the half-shell market (Botta et al. 2020, Summa et al. 2023), and the high prevalence of infestation documented in this study highlights the potential threat these pests pose to the valuable Pacific oyster aquaculture operations in the Mediterranean Sea. The existence of this polydorin, which burrows into oyster shells, represents a threat to the valuable oyster aquaculture industry in the Mediterranean Sea, particularly in the upper Adriatic region where *P. websteri* has currently been located.

Drinkwaard (1998) mentioned that the introduction of *M. gigas*, the Pacific oyster, to northern Holland in 1964 challenged the prevailing assumption that low temperatures would impede its spread. Despite these expectations, reproductive success enabled the establishment of wild populations near the initial introduction sites by the early 1980s. Subsequently, the species expanded its range, reaching the coastlines of Germany and Scandinavia, where it is now recognized as an invasive alien species (Wehrmann et al. 2000). In the Mediterranean Sea, the presence of *M. gigas* is ubiquitous as a consequence of multiple deliberate introductions followed by secondary dispersal. This pattern is exemplified by the Adriatic Sea, where numerous wild specimens have been documented since the 1970s (Šegvić-Bubić et al. 2016). The species was even introduced into the Venice Lagoon as early as 1966 (Marchini et al. 2015).Despite the adaptation and reproductive success of Pacific oysters in Italian waters, oyster farming in Italy remains relatively underdeveloped compared to other European countries. In terms of farming, Italy’s productivity lags behind countries such as Ireland, the Netherlands, Spain, Portugal, and France in terms of productivity. While oyster farming is a well-established industry in other European countries, such as France and the Netherlands, it remains in a nascent stage in Italy (Summa et al. 2023). The Italian oyster farming industry experienced a period of stagnation for many years, until approximately 10-15 years ago when a resurgence began. This resurgence was characterized by the emergence of small-scale production centres scattered across the peninsula. These centres, established in Sardinia, Liguria, Marche, Emilia-Romagna, and Veneto, represent a new wave of independent oyster farming initiatives employing diverse technological approaches. At the moment, they are rather limited experiences (with a total production not exceeding 500 tons per year) and not yet structured as a competitive sector (Summa et al. 2023).

The introduction of *P. websteri* therefore, poses a threat not only to areas of the Mediterranean Sea where oyster farming is already highly developed and productive, but also to the Italian coast, where such an industry is now starting to develop.

The fact that *P. websteri* has been documented in Mediterranean Sea oysters only recently suggests that it was a recent introduction, but it is also possible that the species has been present in the area for some time and has experienced a recent increase in prevalence, which possibly related to expansion of the aquaculture industry or changes in environmental. The expansion of oyster farming in recent years may have led to an increase in the exchange of shell and live oysters between regions and to such an extent that *P. websteri* populations may have further expanded their range and colonized new locations.

In view of the recent discovery of *P. websteri* in the Mediterranean location, it is important to highlight that this spionid polychaete species is known to be sensitive to environmental changes. For instance, rising sediment levels can enhance the vulnerability of *Crassostrea virginica* oysters to *P. websteri* infestation. Conversely, lowering pH levels actually reduces the susceptibility of oysters to this parasitic worm (Clements et al. 2017a, b). Due to its ability to colonize both live and dead oyster shells (Clements et al. 2018), the expansion of the oyster farming, oyster restoration efforts, and the increasing density of oysters in coastal beds may increase the transmission and prevalence of *P. websteri*. Regardless of their origin or extent of exposure to environmental changes, the blister-forming polychaetes documented in this study pose a significant challenge for Mediterranean oyster farmers and shellfish management authorities.

This occurrence highlights the vulnerability of the Adriatic to the invasion of alien species. Notably, many recognized alien species have been found in the upped Adriatic Sea, including molluscs such as *Anadara transversa* (Say, 1822), *Anadara kagoshimensis* (Tokunaga, 1906), *Rapana venosa* (Valenciennes, 1846), *Callinectes sapidus* (Rathbun 1896) (Occhipinti-Ambrogi et al. 2011) e also polychaetes such as *P. cornuta, Chaetozone corona* Berkeley & Berkeley, 1941, *Pseudopolydora paucibranchiata* (Okuda, 1937)(Bertasi 2016, Langeneck et al. 2020, Grossi et al. 2020). This record provides insights into the potential spread of alien species through the oysterfish, specifically in the case of *M. gigas* used in aquaculture. It draws attention to the potential impacts of this species in its new environment, not only affecting the non-native *M. gigas*, but also potentially impacting other native molluscs species. Indeed the generalist nature of *P. websteri* (Surugiu 2005, Read 2010) raises concerns about its potential impact on other shellfish species of ecological, economic, and cultural significance and mussels, scallops and abalone are all endangered extinction (Rodewald et al. 2021). Given the critical ecosystem services provided by filter-feeding shellfish species (van der Schatte Olivier et al. 2020), outbreaks of *P. websteri* could have deleterious consequences beyond the shellfish industry and impact the overall functioning of marine ecosystems.

## References

Abd-Elnaby F. A. 2019. New recorded alien polydorid species (Polychaeta: Spionidae) from the Egyptian waters. Egypt. J. Aquat. Biol. Fish. 23: 409–420.

Bailey-Brock J. H. 1982. Methods for control of the mud blister worm Polydora websteri in Hawaiian oyster culture. Sea grant q. 4: 1–6.

Barros T. L., Gomes Santos C.S., De Assis J. E., Souza J. R. 2017. Morphology and larval development of Polydora cf. websteri (Polychaeta: Spionidae) in a tropical region of north-eastern Brazil. J. Nat. Hist. 51: 1169–1181.

Bertasi F. 2016. The occurrence of the alien species Polydora cornuta Bosc 1802 (Polychaeta: Spionidae) in North Adriatic Lagoons: An overlooked presence. Ital. J. Zool. 83: 77–88.

Blake J. A. 1971. Revision of the genus Polydora from the East Coast of North America (Polychaeta: Spionidae). Smithson. contrib. zool. 1–32.

Blake J. A. 1969. Systematics and ecology of shell-boring polychaetes from New England. Amer. Zool. 9 813–820.

Blake J. A., Kudenov J. D. 1978. The Spionidae (Polychaeta) from southeastern Australia with a revision of the genera of the family. Mem. Mus. Vic. 39: 171–280.

Botta R., Asche F., Borsum J. S., Camp E. V. 2020. A review of global oyster aquaculture production and consumption. Mar. Policy 117: 103952.

Buschbaum C., Buschbaum G., Schrey I., Thieltges D.W. 2007. Shell-boring polychaetes affect gastropod shell strength and crab predation. Mar. Ecol. Prog. Ser., 329: 123–130.

Çinar M. E., Ergen Z., Dagli E., Petersen M. E. 2005. Alien species of Spionid Polychaetes (Streblospio gynobranchiata and Polydora cornuta) in Izmir Bay Eastern Mediterranean. J. Mar. Biol. Assoc. U. K. 85: 821–827.

Clements J. C., Bourque D., McLaughlin J., Stephenson M., Comeau L. A. 2018. Wanted dead or alive: Polydora websteri recruit to both live oysters and empty shells of the Eastern Oyster Crassostrea virginica. J. Fish Dis. 41: 855–858.

Clements J.C., Bourque D., McLaughlin J., Stephenson M., Comeau L.A. 2017a. Siltation Increases the Susceptibility of Surface-cultured Eastern Oysters (Crassostrea virginica) to Parasitism by the Mudworm Polydora websteri. Aquac. Res. 48: 4707–4717.

Clements J.C., Bourque D., McLaughlin J., Stephenson M., Comeau L.A. 2017b. Extreme Ocean Acidification Reduces the Susceptibility of Eastern Oyster Shells to a Polydorid Parasite. J. Fish Dis. 40: 1573–1585.

Coll M., Piroddi C., Steenbeek J., Kaschner K., Ben Rais Lasram F., Aguzzi J., Ballesteros E., Bianchi C. N., Corbera J., Dailianis T., Danovaro R., Estrada M., Froglia C., Galil B. S., Gasol J. M., Gertwagen R., Gil J., Guilhaumon F., Kesner-Reyes K., Kitsos M.-S., Koukouras A., Lampadariou N., Laxamana E., López-Fé de la Cuadra C.M., Lotze H. K., Martin D., Mouillot D., Oro D., Raicevich S., Rius-Barile J., Saiz-Salinas J. I., San Vicente C., Somot S., Templado J., Turon X., Vafidis D., Villanueva R., Voultsiadou E. 2010. The biodiversity of the Mediterranean Sea: Estimates patterns and threats. PLoS ONE 5(8) e11842.

Drinkwaard A. C. 1998. Introductions and Developments of Oysters in the North Sea Area: A Review. Helgoländer Meeresunters. 52 doi:10.1007/BF02908904.

Elton C. S. The Reasons for Conservation. In: The Ecology of Invasions by Animals and Plants. Springer, Boston, MA, London 1958.

Foster N. M. 1971. Spionidae (Polychaeta) of the Gulf of Mexico and the Caribbean Sea. Stud. fauna Curaçao other Caribb. isl. 129: 1–183.

Goedknegt M. A., Feis M. E., Wegner K. M., Luttikhuizen P. C., Buschbaum C., Camphuysen (K.C., van der Meer J., Thieltges D. W. 2016. Parasites and marine invasions: Ecological and evolutionary perspectives. J. Sea Res. 113: 11–27.

Grossi L., Bertasi F., Trabucco B. 2017. New records of the alien polychaete worm Chaetozone corona (Polychaeta: Cirratulidae) in the Adriatic Sea. Acta Adriat. 58: 235–244.

Haigler S. A. 1969. Boring mechanism of Polydora websteri inhabiting Crassostrea virginica. Amer. Zool. 9: 821–828.

Handley S. J., Bergquist P. R. 1997. Spionid polychaete infestations of intertidal pacific oysters Crassostrea gigas (Thunberg) Mahurangi Harbour northern New Zealand. Aquac. 153: 191–205.

Katsanevakis S., Zenetos A., Belchior C., Cardoso A. C. 2013. Invading european seas: Assessing pathways of introduction of marine aliens. Ocean. Coast. Manag. 76: 64–74.

Kent R. M. 1979. The influence of heavy infestations of P. ciliata on the flesh content of Mytilus edulis. J. Mar. Biol. Assoc. U. K. 59: 289–297.

Langeneck J., Lezzi M., Del Pasqua M., Musco L., Gambi M. C., Castelli A., Giangrande A. 2020. Non-indigenous polychaetes along the coasts of Italy: A critical review. Med. Mar. Sci. 2: 238–275.

Leonart M., Handlinger J., Powell M. 2003. Spionid mudworm infestation of farmed abalone (Haliotis spp.). Aquac. 221: 85–96.

Loosanoff V. L., Engle J. B. 1943. Polydora in oysters suspended in the water. Biol. Bull. 85: 69–78.

Lunz G. R. 1940. The annelid worm Polydora as an oyster pest. Science 92: 310–310.

Marchini A., Ferrario J., Sfriso A., Occhipinti-Ambrogi A. 2015. Current Status and Trends of Biological Invasions in the Lagoon of Venice a Hotspot of Marine NIS Introductions in the Mediterranean Sea. Biol. Invasions. 17: 2943–2962.

Martinelli J. C., Lopes H. M., Hauser L., Jimenez-Hidalgo I., King T. L., Padilla-Gamiño J.L., Rawson P., Spencer L. H., Williams J. D., Wood C. L. 2020. Confirmation of the shell-boring oyster parasite Polydora websteri (Polychaeta: Spionidae) in Washington State USA. Sci. Rep. 10: 3961.

Mckindsey C. W., Landry T., O’Beirn F. X., Davies I. M. 2007. Bivalve aquaculture and exotic species: A review of ecological considerations and management issues. J. Shellfish Res. 26: 281–294.

Mikac B., Fossi E., Costantini F., Colangelo M.A., Tarullo A., Prioli G., Radashevsky V.I. 2023. New alien Polydora oysters; pests in the Mediterranean. Programme Abstracts of the 14th International Polychaete Conference. Stellenbosch: Stellenbosch University. P. 75–76.

Morse D. L., Rawson P. D., Kraeuter J. N. 2015. Mud blister worms and oyster aquaculture. Maine Sea Grant Publications. Available at: https://digitalcommons.library.umaine.edu/seagrant_pub/46/

Occhipinti-Ambrogi A., Marchini A., Cantone G., Castelli A., Chimenz C., Cormaci M., Froglia C., Furnari G., Gambi M. C., Giaccone G., Giangrande A., Gravili C., Mastrototaro F., Mazziotti C., Orsi-Relini L., Piraino S. 2010. Alien species along the Italian coasts: An overview. Biol. Invasions 13: 215–237.

Ogburn D. M., White I., Mcphee D. P. 2007. The disappearance of oyster reefs from eastern Australian estuaries - impact of colonial settlement or mudworm invasion? Coast. Manage. 35: 271–287.

Padilla D. K., McCann M. J., Shumway S. E. 2011. Marine invaders and bivalve aquaculture: Sources impacts and consequences. In Shellfish Aquaculture and the Environment ed. by Shumway SE. Wiley Blackwell Hoboken NY

Radashevsky V. I. 1999. Description of the proposed lectotype for Polydora websteri Hhartman in Loosanoff & Engle 1943 (Polychaeta: Spionidae). Ophelia 51: 107–113.

Radashevsky V. I. 2005. On adult and larval morphology of Polydora cornuta Bosc 1802 (Annelida: Spionidae). Zootaxa 1064: 1–24.

Radashevsky V. I., Williams J. D. 1998. Polydora websteri Hartman in Loosanoff & Engle 1943 (Annelida Polychaeta): Proposed conservation of the specific name by a ruling that it is not to be treated as a replacement for P. caeca Webster 1879 and designation of a lectotype for P. websteri. Bull. zool. nomencl. 55: 212–216.

Read G. B. 2010. Comparison and history of Polydora websteri and P. haswelli (Polychaeta: Spionidae) as mud-blister worms in New Zealand shellfish. N. Z. J. Mar. Freshwater Res. 44: 83–100.

Rice L., Lindsay S., Rawson P. 2018. Genetic homogeneity among geographically distant populations of the blister worm Polydora websteri. Aquac. Environ. Interact. 10: 437–446.

Rodewald N., Snyman R., Simon C. A. 2021. Worming its way in Polydora websteri (Annelida: Spionidae) increases the number of non-indigenous shell-boring polydorin pests of cultured molluscs in South Africa. Zootaxa 4969.

Ruesink J. L., Lenihan H. S., Trimble A. C., Heiman K. W., Micheli F., Byers J. E., Kay M. C. 2005. Introduction of non-native oysters: Ecosystem effects and restoration implications. Annu. Rev. Ecol. Evol. Syst. 36: 643–689.

Sato-Okoshi W. 1999. Polydorid species (Polychaeta: Spionidae) in Japan with descriptions of morphology ecology and burrow structure. 1. Boring species. J. Mar. Biol. Assoc. U. K. 79: 831–848.

Sato-Okoshi W., Abe H. 2013. Morphology and molecular analysis of the 18S r-RNA gene of oyster shell borers Polydora species (Polychaeta: Spionidae) from Japan and Australia. J. Mar. Biol. Assoc. U. K. 93: 1279–1286.

Šegvić-Bubić T., Grubišić L., Zrnčić S., Jozić S., žužul I., Talijančić I., Oraić D., Relić M., Katavić I. 2016. Range Expansion of the Non-native Oyster Crassostrea Gigas in the Adriatic Sea. Acta Adriat. 57: Polydora websteri can be distinguished 142: 196-270.

Simon C., Sato-Okoshi W. 2015. Polydorid polychaetes on farmed molluscs: Distribution spread and factors contributing to their success. Aquac. Environ. Interact. 7: 147–166.

Summa D. M., Turolla E, Lanzoni M, Tamisari E, Castaldelli G, Tamburini E. 2023. Life Cycle Assessment (LCA) of Two Different Oyster (Crassostrea gigas) farming Strategies in the Sacca di Goro Northern Adriatic Sea Italy. Resources. 12(6): 62.

Surugiu V. 2005. Inventory of inshore polychaetes from the Romanian coast (Black Sea). Med. Mar. Sci. 6(1): 51–74.

Surugiu V. 2012. Systematics and ecology of species of the Polydora-complex (Polychaeta: Spionidae) of the Black Sea. Zootaxa 3518: 45.

Tan K., Cheng D., Kwan K. Y., Peng Y., Cai X., Lim L., Xu P., Tan K. 2023. Research progress of shell boring mud-blister worm infestation in shellfish aquaculture. Aquac. 574: 739–693.

van der Schatte Olivier A., Jones L., Le Vay L., Christie M., Wilson J.M., Malham S.K. 2020. A Global Review of the Ecosystem Services Provided by Bivalve Aquaculture. Rev. Aquac. 12: 3–25.

Waser A. M., Lackschewitz D., Knol J., Reise K., Wegner K. M., Thieltges D. W. 2020. Spread of the invasive shell-boring annelid Polydora websteri (Polychaeta Spionidae) into naturalized oyster reefs in the European Wadden Sea. Mar. Biodivers. 50.

Wehrmann A., Herlyn M., Bungenstock F., Hertweck G., Millat G. 2000. The Distribution Gap Is Closed - First Record of Naturally Settled Pacific Oysters Crassostrea Gigas in the East Frisian Wadden Sea North Sea. Mar. Biodivers. 30 doi:10.1007/BF03042964.

Whitelegge T. 1890. Report on the worm disease affecting the oysters on the coast of New South Wales. Rec. Aust. Mus. 1:41–54.

Wolff W. J., Reise K. 2002. Oyster imports as a vector for the introduction of alien species into northern and Western European coastal waters. Invasive Aquatic Species of Europe. Distribution Impacts and Management 193–205.

Ye L., Cao C., Tang B., Yao T., Wang R., Wang J. 2017. Morphological and molecular characterization of Polydora websteri (Annelida: Spionidae) with remarks on relationship of adult worms and larvae using mitochondrial COI gene as a molecular marker. Pak. J. Zool. 49: 699–710.

Zottoli R. A., Carriker M. R. 1974. Burrow morphology tube formation and microarchitecture of shell dissolution by the Spionid Polychaete Polydora websteri. Mar. Biol. 27: 307–316.

